# Comparative Genomic Analysis of *Salmonella enterica* serovar Typhimurium from Passerines Reveals Two Lineages Circulating in Europe, New Zealand, and the United States

**DOI:** 10.1101/2022.03.08.483506

**Authors:** Yezhi Fu, Nkuchia M. M’ikanatha, Edward G. Dudley

## Abstract

*Salmonella enterica* serovar Typhimurium from passerines have caused wild bird mortality and human salmonellosis outbreaks in Europe, Oceania, and North America. Here, we performed comparative genomic analysis to explore the emergence, genetic relationship, and evolution of geographically dispersed passerine isolates. We found that passerine isolates from Europe and the United States clustered to form two lineages (EU and US passerine lineages), which were distinct from major *S*. Typhimurium lineages circulating in other diverse hosts (*e.g*., humans, cattle, pigs, chicken, other avian hosts such as pigeons and ducks). Further, passerine isolates from New Zealand clustered to form a sublineage (NZ passerine lineage) of the US passerine lineage. We inferred that the passerine isolates mutated at a rate of 3.2 × 10^-7^ substitutions/site/year, and the US, EU, and NZ passerine lineages emerged in *ca*. 1952, 1970, and 1996, respectively. Isolates from the three lineages presented genetic similarity such as lack of antimicrobial resistance genes and accumulation of same virulence pseudogenes. In addition, genetic diversity due to microevolution existed in the three passerine lineages. Specifically, pseudogenization in type 1 fimbrial gene *fimC* (deletion of G at position 87) was only detected in the US and NZ passerine isolates, while a single-base deletion in type 3 secretion system effector genes (*i.e*., *gogB, sseJ*, and *sseK2*) solely concurred in the EU passerine isolates. These findings provide insights into evolution, host adaptation, and epidemiology of *S*. Typhimurium in passerines.

**IMPORTANCE:** Passerine-associated *S*. Typhimurium have been linked to human salmonellosis outbreaks in recent years. Here we investigated the phylogenetic relationship of globally distributed passerine isolates and profiled their genomic similarity and diversity. Our study reveals two passerine-associated *S*. Typhimurium lineages circulating in Europe, Oceania, and North America. Isolates from the two lineages presented phylogenetic and genetic signatures that were distinct from isolates of other hosts. The findings shed light on host adaptation of *S*. Typhimurium in passerines and are important for source attribution of *S*. Typhimurium to avian hosts. Further, we found *S*. Typhimurium definitive phage type (DT) 160 from passerines that caused decade-long human salmonellosis outbreaks in New Zealand and Australia formed a sublineage of the US passerine lineage, suggesting that DT160 may have originated from passerines outside Oceania. Our study demonstrates the importance of whole-genome sequencing and genomic analysis of historical microbial collections to modern day epidemiologic surveillance.

## INTRODUCTION

*Salmonella enterica* serovar Typhimurium is a leading cause of salmonellosis worldwide. Generally, *S*. Typhimurium can colonize and infect a broad range of hosts such as humans, livestock, poultry, and wild animals. Examples of broad-host-range *S*. Typhimurium variants circulating worldwide include *S*. Typhimurium definitive phage type (DT) 104 (1) and monophasic *S*. Typhimurium (*S*. 4,[5],12:i:-) sequence type (ST) 34 (2). However, some variants of *S*. Typhimurium are subjected to continuous evolution within specific hosts, thus exhibiting host preference or adaptation. These variants are primarily found in wild birds, which include *S*. Typhimurium DT2 and DT99 from feral pigeons (3), DT8 linked to ducks (4), and DT40 and DT56(v) associated with passerine birds (*i.e*., any birds in the order *Passeriformes* such as sparrows, siskins, finches) (5, 6). Although the above-mentioned *S*. Typhimurium variants have host tropism to particular wild birds, they occasionally infect humans, domestic animals, and other host species.

In the past decades, passerine-associated *S*. Typhimurium have been linked to salmonellosis outbreaks in both humans and wild birds. In New Zealand, a decade-long (1998-2012) outbreak of *S*. Typhimurium DT160 affected >3,000 people and killed passerines (7). In 2008, *S*. Typhimurium DT160 was identified in Tasmania, Australia, where it infected ≈ 50 people and caused passerine mortality (8). In Europe, outbreaks of passerine-associated human infections have been reported in the United Kingdom (5, 9) and Sweden (10). These outbreaks were caused by *S*. Typhimurium DT40 and DT56(v). In the United States, *S*. Typhimurium isolates associated with the 2009 pine siskin outbreak were implicated in a human salmonellosis outbreak in the same year (11). More recently, a 2021 *S*. Typhimurium outbreak linked to passerines resulted in 29 illnesses and 14 hospitalizations in 12 US states (12).

The emergence of passerine-associated *S*. Typhimurium and corresponding outbreaks worldwide raises the questions about their origin, evolution, and genetic relationship. In this study, we conducted comparative genomic analysis of passerine-associated *S*. Typhimurium from Europe, New Zealand, and the United States over a 40-year period. The genetic relationship and emergence of passerine-associated *S*. Typhimurium from different locations were inferred by phylogenetic analysis and Bayesian inference, respectively. Further, we investigated the genetic content of passerine-associated *S*. Typhimurium by profiling their virulence factors, plasmids, and antimicrobial resistance (AMR) determinants. We also compared the whole-genome sequences of *S*. Typhimurium from passerines and other diverse hosts (*e.g*., humans, cattle, pigs, poultry, other birds such as pigeons and ducks) to determine if passerine-associated *S*. Typhimurium had distinct phylogenetic and genetic signatures.

## RESULTS

### Phylogenetic relationship of geographically dispersed *S*. Typhimurium from passerines

A maximum-likelihood phylogenetic tree (Figure 1) was built based on 2,253 single nucleotide polymorphisms (SNPs) in the core genomic regions of 84 publicly available passerine isolates (Table 1; New Zealand: *n* = 25, isolated year: 2000–2009; United States: *n* = 33, isolated year: 1978–2019; European countries: United Kingdom: *n* = 11, Sweden: *n* = 14, Germany: *n* = 1, isolated year: 2001–2016) against reference genome of *S*. Typhimurium strain LT2 (RefSeq NC_003197.1). We found that passerine isolates from Europe clustered to form a lineage (henceforth referred to as the EU passerine lineage). Further, we observed that passerine isolates from the United States clustered to form a lineage (henceforth referred to as the US passerine lineage), and *S*. Typhimurium DT160 isolates from New Zealand clustered to form a sublineage (henceforth referred to as the NZ passerine lineage) of the US passerine lineage (Figure 1). The EU, US, and NZ passerine lineages were supported by robust bootstrap values of 100%. The average SNP distance in the core genome between isolates in the US and NZ passerine lineages was 81, while the average SNP distance in the core genome between isolates in the US, NZ, and EU passerine lineages was 265. Multilocus sequence typing (MLST) indicated that isolates from NZ and US passerine lineages belonged to ST19, whereas the European passerine isolates presented variable STs (*i.e*., ST19, 568, and 7075).

**Table 1.** Metadata information of the 84 *Salmonella enterica* serovar Typhimurium isolates from passerines.

| Isolate name | Collection year | Continent | Country | Accession number | Source | Source detail | Sequence type |
| --- | --- | --- | --- | --- | --- | --- | --- |
| 054/01 | 2001 | Europe | United Kingdom | ERS217365 | Passeriformes | House sparrow | 568 |
| 062/01 | 2001 | Europe | United Kingdom | ERS217362 | Passeriformes | European greenfinch | 19 |
| 065/01 | 2001 | Europe | United Kingdom | ERS217360 | Passeriformes | House sparrow | 19 |
| 100/01 | 2001 | Europe | United Kingdom | ERS217364 | Passeriformes | European greenfinch | 19 |
| 108/01 | 2001 | Europe | United Kingdom | ERS217358 | Passeriformes | European greenfinch | 568 |
| 132/06 | 2006 | Europe | United Kingdom | ERS217361 | Passeriformes | European greenfinch | 568 |
| 1356/06 | 2006 | Europe | United Kingdom | ERS217366 | Passeriformes | House sparrow | 19 |
| 1377/06 | 2006 | Europe | United Kingdom | ERS217363 | Passeriformes | House sparrow | 568 |
| 1402/06 | 2006 | Europe | United Kingdom | ERS217356 | Passeriformes | European greenfinch | 19 |
| 1422/05 | 2005 | Europe | United Kingdom | ERS217359 | Passeriformes | European greenfinch | 568 |
| 20-SA01024-0 | 2020 | Europe | Germany | SRS9340518 | Passeriformes | Eurasian blue tit | 19 |
| S00228-16 | 2016 | Europe | United Kingdom | ERS2028946 | Passeriformes | Finch | 19 |
| STm SE 01 | 2016 | Europe | Sweden | ERR2617828 | Passeriformes | Common redpoll | 19 |
| STm SE 02 | 2016 | Europe | Sweden | ERR2617829 | Passeriformes | Common redpoll | 19 |
| STm SE 03 | 2016 | Europe | Sweden | ERR2617830 | Passeriformes | Common redpoll | 19 |
| STm SE 04 | 2016 | Europe | Sweden | ERR2617831 | Passeriformes | Common redpoll | 19 |
| STm SE 05 | 2016 | Europe | Sweden | ERR2617832 | Passeriformes | Eurasian bullfinch | 19 |
| STm SE 06 | 2016 | Europe | Sweden | ERR2617833 | Passeriformes | Eurasian bullfinch | 19 |
| STm SE 08 | 2016 | Europe | Sweden | ERR2617835 | Passeriformes | Eurasian bullfinch | 7075 |
| STm SE 09 | 2016 | Europe | Sweden | ERR2617836 | Passeriformes | Eurasian siskin | 19 |
| STm SE 10 | 2016 | Europe | Sweden | ERR2617837 | Passeriformes | Eurasian siskin | 19 |
| STm SE 11 | 2016 | Europe | Sweden | ERR2617838 | Passeriformes | Eurasian bullfinch | 19 |
| STm SE 12 | 2016 | Europe | Sweden | ERR2617839 | Passeriformes | Eurasian bullfinch | 19 |
| STm SE 13 | 2016 | Europe | Sweden | ERR2617840 | Passeriformes | Eurasian bullfinch | 19 |
| STm SE 14 | 2016 | Europe | Sweden | ERR2617841 | Passeriformes | Eurasian bullfinch | 19 |
| STm SE 15 | 2016 | Europe | Sweden | ERR2617842 | Passeriformes | Eurasian bullfinch | 19 |
| PSU-2812 | 1998 | North America | United States | SRS7318189 | Passeriformes | Evening grosbeak | 19 |
| PSU-2813 | 1998 | North America | United States | SRS7318190 | Passeriformes | Redpoll | 19 |
| PSU-2814 | 1998 | North America | United States | SRS7318191 | Passeriformes | American goldfinch | 19 |
| PSU-2816 | 1998 | North America | United States | SRS7318193 | Passeriformes | White-throated sparrow | 19 |
| PSU-2817 | 1998 | North America | United States | SRS7417389 | Passeriformes | Redpoll | 19 |
| PSU-2819 | 2000 | North America | United States | SRS7318196 | Passeriformes | Gold finch | 19 |
| PSU-2835 | 2009 | North America | United States | SRS7611982 | Passeriformes | Purple finch | 19 |
| PSU-2838 | 2009 | North America | United States | SRS7611983 | Passeriformes | Pine siskin | 19 |
| PSU-2841 | 2009 | North America | United States | SRS7417441 | Passeriformes | House sparrow | 19 |
| PSU-2847 | 2011 | North America | United States | SRS7417395 | Passeriformes | House sparrow | 19 |
| PSU-2966 | 1978 | North America | United States | SRS7417415 | Passeriformes | House sparrow | 19 |
| PSU-3168 | 1978 | North America | United States | SRS7449455 | Passeriformes | House sparrow | 19 |
| PSU-3169 | 1979 | North America | United States | SRS7718306 | Passeriformes | House sparrow | 19 |
| PSU-3174 | 1991 | North America | United States | SRS7449460 | Passeriformes | Goldfinch | 19 |
| PSU-3234 | 1992 | North America | United States | SRS7612006 | Passeriformes | Cardinal | 19 |
| PSU-3235 | 1992 | North America | United States | SRS7718330 | Passeriformes | House sparrow | 19 |
| PSU-3236 | 1992 | North America | United States | SRS7612008 | Passeriformes | House sparrow | 19 |
| PSU-3253 | 1991 | North America | United States | SRS7611956 | Passeriformes | Brown-headed cowbird | 19 |
| PSU-3337 | 2018 | North America | United States | SRS7840968 | Passeriformes | Redpoll | 19 |
| PSU-3338 | 2019 | North America | United States | SRS7840979 | Passeriformes | Pine siskin | 19 |
| PSU-3340 | 2016 | North America | United States | SRS7840996 | Passeriformes | House sparrow | 19 |
| PSU-3346 | 2016 | North America | United States | SRS7840969 | Passeriformes | Red-winged blackbird | 19 |
| PSU-3353 | 2015 | North America | United States | SRS7840976 | Passeriformes | Red crossbill | 19 |
| PSU-3358 | 2016 | North America | United States | SRS7840981 | Passeriformes | Pine siskin | 19 |
| PSU-3359 | 2015 | North America | United States | SRS7840982 | Passeriformes | Red crossbill | 19 |
| PSU-3363 | 2016 | North America | United States | SRS7840986 | Passeriformes | Redpoll | 19 |
| PSU-3367 | 2018 | North America | United States | SRS7840991 | Passeriformes | Pine siskin | 19 |
| PSU-3374 | 2018 | North America | United States | SRS7841677 | Passeriformes | Redpoll | 19 |
| PSU-3377 | 2016 | North America | United States | SRS7841693 | Passeriformes | Pine siskin | 19 |
| PSU-3378 | 2015 | North America | United States | SRS7841694 | Passeriformes | Red crossbill | 19 |
| PSU-3379 | 2015 | North America | United States | SRS7841695 | Passeriformes | Red crossbill | 19 |
| PSU-3405 | 2013 | North America | United States | SRS7841689 | Passeriformes | Pine siskin | 19 |
| PSU-4760 | 2012 | North America | United States | SRS9461411 | Passeriformes | American goldfinch | 19 |
| DT160 01 | 2000 | Oceania | New Zealand | ERS1456804 | Passeriformes | Passerine bird, not specified | 19 |
| DT160 02 | 2007 | Oceania | New Zealand | ERS1456799 | Passeriformes | Passerine bird, not specified | 19 |
| DT160 03 | 2000 | Oceania | New Zealand | ERS1456794 | Passeriformes | Passerine bird, not specified | 19 |
| DT160 04 | 2006 | Oceania | New Zealand | ERS1456793 | Passeriformes | Passerine bird, not specified | 19 |
| DT160 05 | 2008 | Oceania | New Zealand | ERS1456792 | Passeriformes | Passerine bird, not specified | 19 |
| DT160 06 | 2003 | Oceania | New Zealand | ERS1456791 | Passeriformes | Passerine bird, not specified | 19 |
| DT160 07 | 2004 | Oceania | New Zealand | ERS1456790 | Passeriformes | Passerine bird, not specified | 19 |
| DT160 08 | 2003 | Oceania | New Zealand | ERS1456787 | Passeriformes | Passerine bird, not specified | 19 |
| DT160 09 | 2007 | Oceania | New Zealand | ERS1456781 | Passeriformes | Passerine bird, not specified | 19 |
| DT160 10 | 2003 | Oceania | New Zealand | ERS1456778 | Passeriformes | Passerine bird, not specified | 19 |
| DT160 11 | 2005 | Oceania | New Zealand | ERS1456773 | Passeriformes | Passerine bird, not specified | 19 |
| DT160 12 | 2001 | Oceania | New Zealand | ERS1456770 | Passeriformes | Passerine bird, not specified | 19 |
| DT160 13 | 2007 | Oceania | New Zealand | ERS1456766 | Passeriformes | Passerine bird, not specified | 19 |
| DT160 14 | 2009 | Oceania | New Zealand | ERS1456765 | Passeriformes | Passerine bird, not specified | 19 |
| DT160 15 | 2008 | Oceania | New Zealand | ERS1456760 | Passeriformes | Passerine bird, not specified | 19 |
| DT160 16 | 2002 | Oceania | New Zealand | ERS1456758 | Passeriformes | Passerine bird, not specified | 19 |
| DT160 17 | 2009 | Oceania | New Zealand | ERS1456757 | Passeriformes | Passerine bird, not specified | 19 |
| DT160 18 | 2004 | Oceania | New Zealand | ERS1456756 | Passeriformes | Passerine bird, not specified | 19 |
| DT160 19 | 2002 | Oceania | New Zealand | ERS1456755 | Passeriformes | Passerine bird, not specified | 19 |
| DT160 20 | 2002 | Oceania | New Zealand | ERS1456754 | Passeriformes | Passerine bird, not specified | 19 |
| DT160 21 | 2001 | Oceania | New Zealand | ERS1456749 | Passeriformes | Passerine bird, not specified | 19 |
| DT160 22 | 2001 | Oceania | New Zealand | ERS1456748 | Passeriformes | Passerine bird, not specified | 19 |
| DT160 23 | 2005 | Oceania | New Zealand | ERS1456744 | Passeriformes | Passerine bird, not specified | 19 |
| DT160 24 | 2005 | Oceania | New Zealand | ERS1456734 | Passeriformes | Passerine bird, not specified | 19 |
| DT160 25 | 2006 | Oceania | New Zealand | ERS1456733 | Passeriformes | Passerine bird, not specified | 19 |

**Figure 1.**
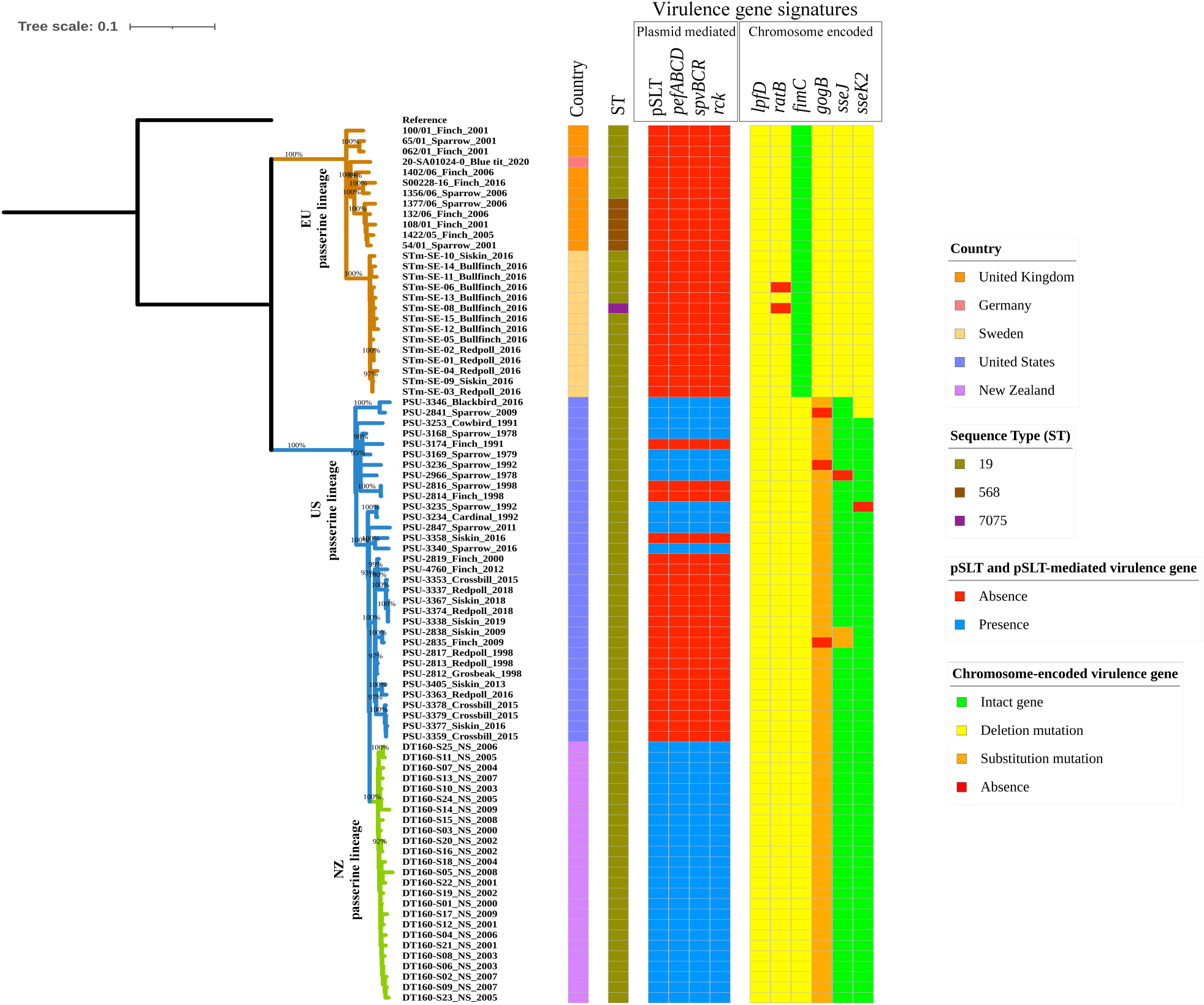
Maximum-likelihood phylogenetic tree of the 84 *Salmonella enterica* serovar Typhimurium isolates from passerines (New Zealand: *n* = 25, United States: *n* = 33, United Kingdom: *n* = 11, Sweden: *n* = 14, Germany: *n* = 1). The tree is created based on 2,253 core-genome single nucleotide polymorphisms (SNPs) with reference to *S*. Typhimurium LT2. Three lineages are defined in the tree, *i.e*., EU (European countries) passerine lineage (brown), US (the United States) passerine lineage (blue), and NZ (New Zealand) passerine lineage (green). Bootstrap values are displayed as percentage on the tree branches. The labels at the tree tips represent the “isolate name_bird host_collection year”. “NS” is used to replace “bird host” if the bird host is not specified in the NCBI database. The color strip “country” represents the isolation location. The color strip “ST” represents the *S*. Typhimurium multilocus sequence type. The virulence gene signatures identified in this study are categorized into plasmid-mediated and chromosome-encoded, and represented in different color strips.

### Emergence time of passerine-associated *S*. Typhimurium lineages in Europe, New Zealand, and the United States

A time-scaled Bayesian phylogenetic tree was built using BEAST2 (v2.6.5) to infer the emergence time of the passerine lineages (Figure 2). The most recent common ancestor (MRCA) of the passerine isolates was estimated to originate in *ca*. 1840 [95% highest probability density (HPD): 1784–1887]. Based on the Bayesian inference, the MRCA evolved to form the US and EU passerine lineages in *ca*. 1952 (95% HPD: 1942–1960) and *ca*. 1970 (95% HPD: 1960–1978), respectively (Figure 2). The NZ passerine lineage formed two sublineages, which emerged in *ca*. 1995 (95% HPD: 1992–1997) and *ca*. 1997 (95% HPD: 1994– 1999) (Figure 2). We estimated that the median substitution rate for the 84 passerine isolates was 3.2 × 10^-7^ substitutions/site/year (95% HPD: 1.8–5.0 × 10^-7^ substitutions/site/year). Median substitution rates for the isolates from the EU and US passerine lineages were 3.2 × 10^-7^ substitutions/site/year (95% HPD: 1.8–4.6 × 10^-7^ substitutions/site/year) and 3.6 × 10^-7^ substitutions/site/year (95% HPD: 2.3–5.5 × 10^-7^ substitutions/site/year), respectively. The isolates from the NZ passerine lineage mutated at the same rate as the US passerine isolates.

**Figure 2.**
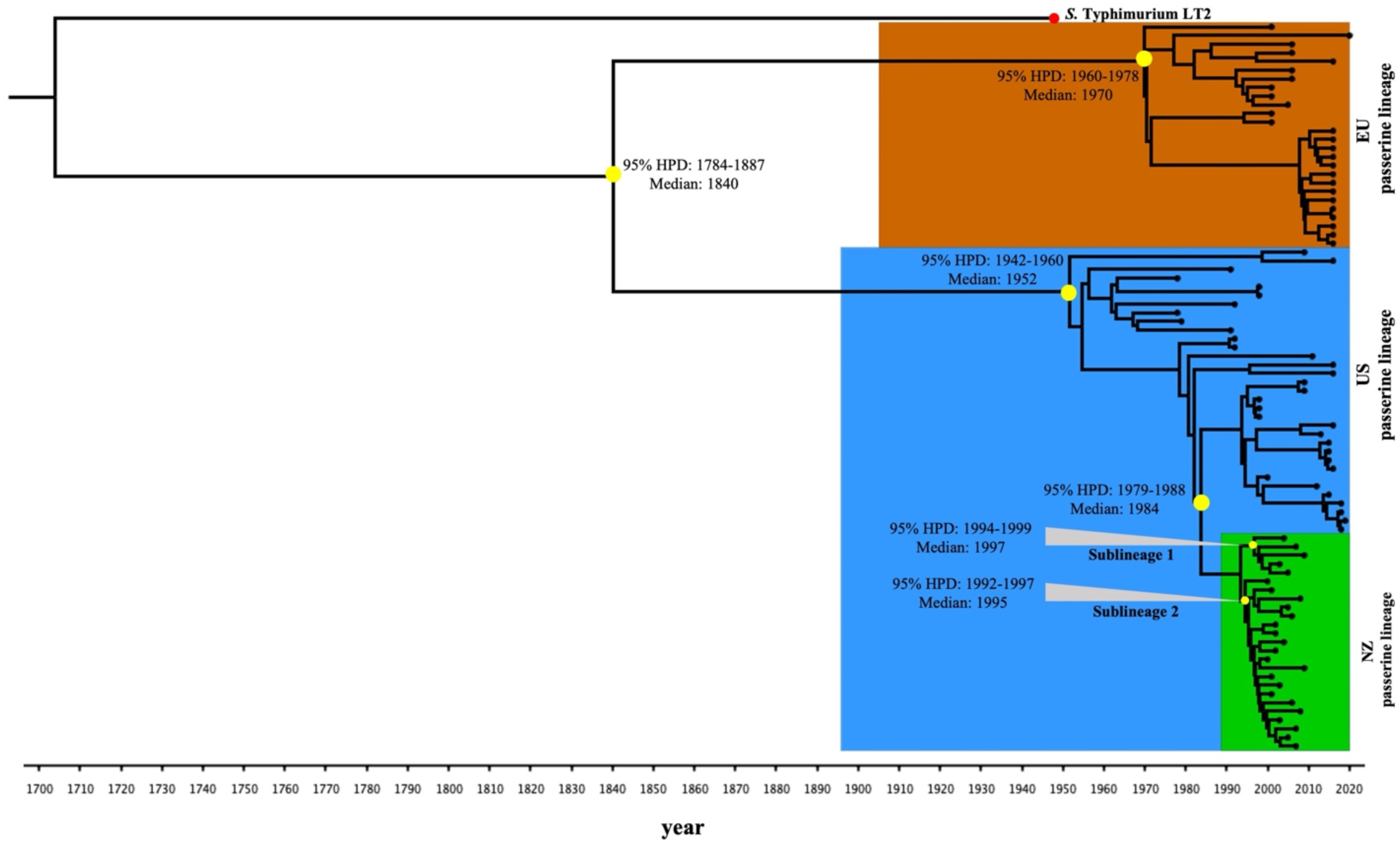
Time-scaled Bayesian phylogenetic tree of the 84 *Salmonella enterica* serovar Typhimurium isolates from passerines. The EU, US, and NZ passerine lineages are colored in brown, blue, and green in the tree, respectively. Median years or range of years at the tree nodes (yellow circles) represent the 95% highest posterior probability density (HPD) for the times of most recent common ancestor for representative divergent events. The red circle at the tree tip represents the reference strain LT2 (collection year: *ca*.1948). The posterior probability values of representative divergent events (yellow circles at tree nodes) are >95%.

### Antimicrobial resistance, plasmid, and virulence gene profiles of passerine-associated *S*. Typhimurium

Antimicrobial resistance (AMR) profiling (Dataset S1) by ResFinder 2.0 detected no AMR genes in isolates from the EU, US, and NZ passerine lineages, except the unexpressed gene *aac(6’)-Iaa* (13). Plasmid profiling (Dataset S1) by PlasmidFinder 2.0 suggested that all of the EU passerine isolates (26/26) lacked the *S*. Typhimurium-specific virulence plasmid pSLT (Figure 1). However, all of the passerine isolates (25/25) from New Zealand and one third (11/33) of the US passerine isolates carried this plasmid (Figure 1). Virulence gene profiling by ABRicate against the Virulence Factor Database (VFDB) database detected an average number of 107, 110, and 116 virulence genes in the EU, US, and NZ passerine isolates, respectively (Dataset S1). The absent virulence genes in the EU and US passerine isolates were primarily plasmid mediated [*i.e*., pSLT-mediated virulence genes: *pefABCD* (plasmid-encoded fimbriae), *rck* (resistance to complement killing), and *spvBCR* (*Salmonella* plasmid virulence)] (Dataset S1; Figure 1). Isolates from the EU, US, and NZ passerine lineages possessed the same chromosomal pseudogenes (*i.e*., *lpfD* and *ratB*) (Figure 1; Table 2). In addition, isolates from the NZ and US passerine lineages had a single-base deletion mutation in type 1 fimbrial gene *fimC*, which was intact in the EU passerine isolates (Figure 1; Table 2). In contrast, single-base deletion mutation in type 3 secretion system (T3SS) effector genes (*i.e*., *gogB, sseJ*, and *sseK2*) was detected in all of the European passerine isolates, however, most of the US and NZ passerine isolates had a single-base substitution rather than deletion mutation in the *gogB* gene, and their *sseJ* and *sseK2* genes were intact (Figure 1; Table 2).

**Table 2.**
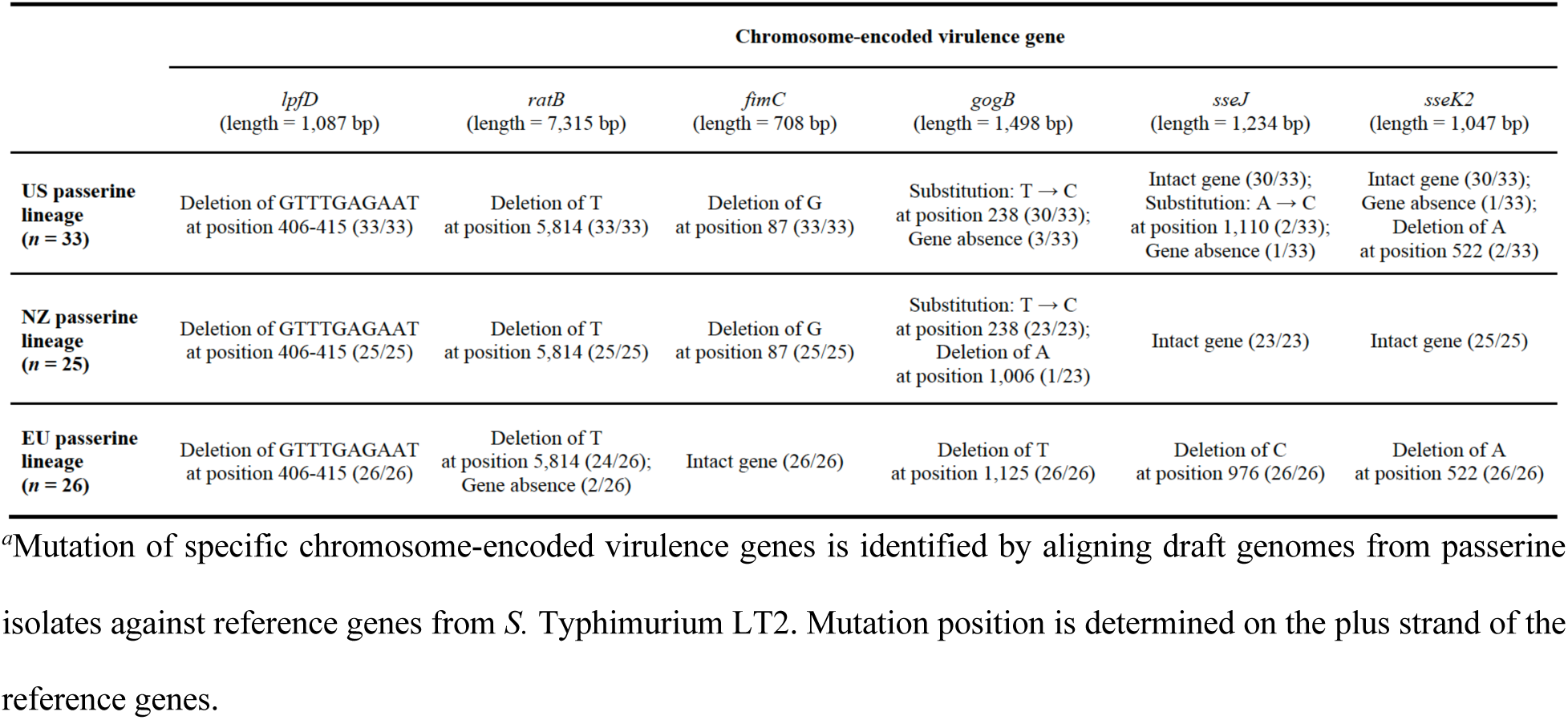
Mutation of specific chromosome-encoded virulence genes in *Salmonella enterica* serovar Typhimurium isolates from passerines^*a*^.

|  | Chromosome-encoded virulence gene |  |  |  |  |  |
| --- | --- | --- | --- | --- | --- | --- |
|  | *lpfD*<br>(length = 1,087 bp) | *ratB*<br>(length = 7,315 bp) | *fimC*<br>(length = 708 bp) | *gogB*<br>(length = 1,498 bp) | *sseJ*<br>(length = 1,234 bp) | *sseK2*<br>(length = 1,047 bp) |
| **US passerine lineage**<br>(*n* = 33) | Deletion of GTTTGAGAAT at position 406-415 (33/33) | Deletion of T at position 5,814 (33/33) | Deletion of G at position 87 (33/33) | Substitution: T → C at position 238 (30/33);<br>Gene absence (3/33) | Intact gene (30/33);<br>Substitution: A → C at position 1,110 (2/33);<br>Gene absence (1/33) | Intact gene (30/33);<br>Gene absence (1/33);<br>Deletion of A at position 522 (2/33) |
| **NZ passerine lineage**<br>(*n* = 25) | Deletion of GTTTGAGAAT at position 406-415 (25/25) | Deletion of T at position 5,814 (25/25) | Deletion of G at position 87 (25/25) | Substitution: T → C at position 238 (23/23);<br>Deletion of A at position 1,006 (1/23) | Intact gene (23/23) | Intact gene (25/25) |
| **EU passerine lineage**<br>(*n* = 26) | Deletion of GTTTGAGAAT at position 406-415 (26/26) | Deletion of T at position 5,814 (24/26);<br>Gene absence (2/26) | Intact gene (26/26) | Deletion of T at position 1,125 (26/26) | Deletion of C at position 976 (26/26) | Deletion of A at position 522 (26/26) |

<sup>a</sup>Mutation of specific chromosome-encoded virulence genes is identified by aligning draft genomes from passerine isolates against reference genes from *S. Typhimurium* LT2. Mutation position is determined on the plus strand of the reference genes.

### Population structure of *S*. Typhimurium from passerines and other diverse hosts

A maximum-likelihood phylogenetic tree (Figure 3A) was built based on 10,065 SNPs in the core genomic regions of passerine isolates (*n* = 84) and context isolates (*n* = 112; Dataset S2) from multiple hosts to represent a broader collection of *S*. Typhimurium. The context isolates formed nine context lineages in the tree (Figure 3A). Six out of the nine context lineages were associated with specific hosts, *i.e*., DT2 (*n* = 13) and DT99 (*n* = 6) lineages adapted to pigeons, DT8 lineage (*n* = 10) associated with ducks, ST313 lineage (*n* = 10) causing invasive nontyphoidal *Salmonella* diseases in humans, DT204 complex lineage (*n* = 9) primarily infecting cattle, and U288 complex lineage (*n* = 20) majorly found in pigs. Additionally, three out of the nine context lineages had broad host range, which included DT104 complex (*n* = 14), DT193 complex (*n* = 9), and ST34 (*n* = 21). We found that the EU (*n* = 26), US (*n* = 33), and NZ (*n* = 25) passerine lineages clustered in a large lineage that was distinct from the nine major *S*. Typhimurium lineages circulating globally in different hosts (Figure 3A). The three passerine lineages had the closest genetic relatedness with DT204 complex lineage (primary host: cattle) in the tree (average SNP distance in the core genome ≈ 308). We also generated a neighbor joining (NJ) tree (Figure 3B) of the 84 passerine and 112 context isolates based on EnteroBase whole-genome MLST. The lineages present in the NJ tree were congruent with those formed in the maximum-likelihood phylogenetic tree based on SNPs (Figure 3B).

**Figure 3.**
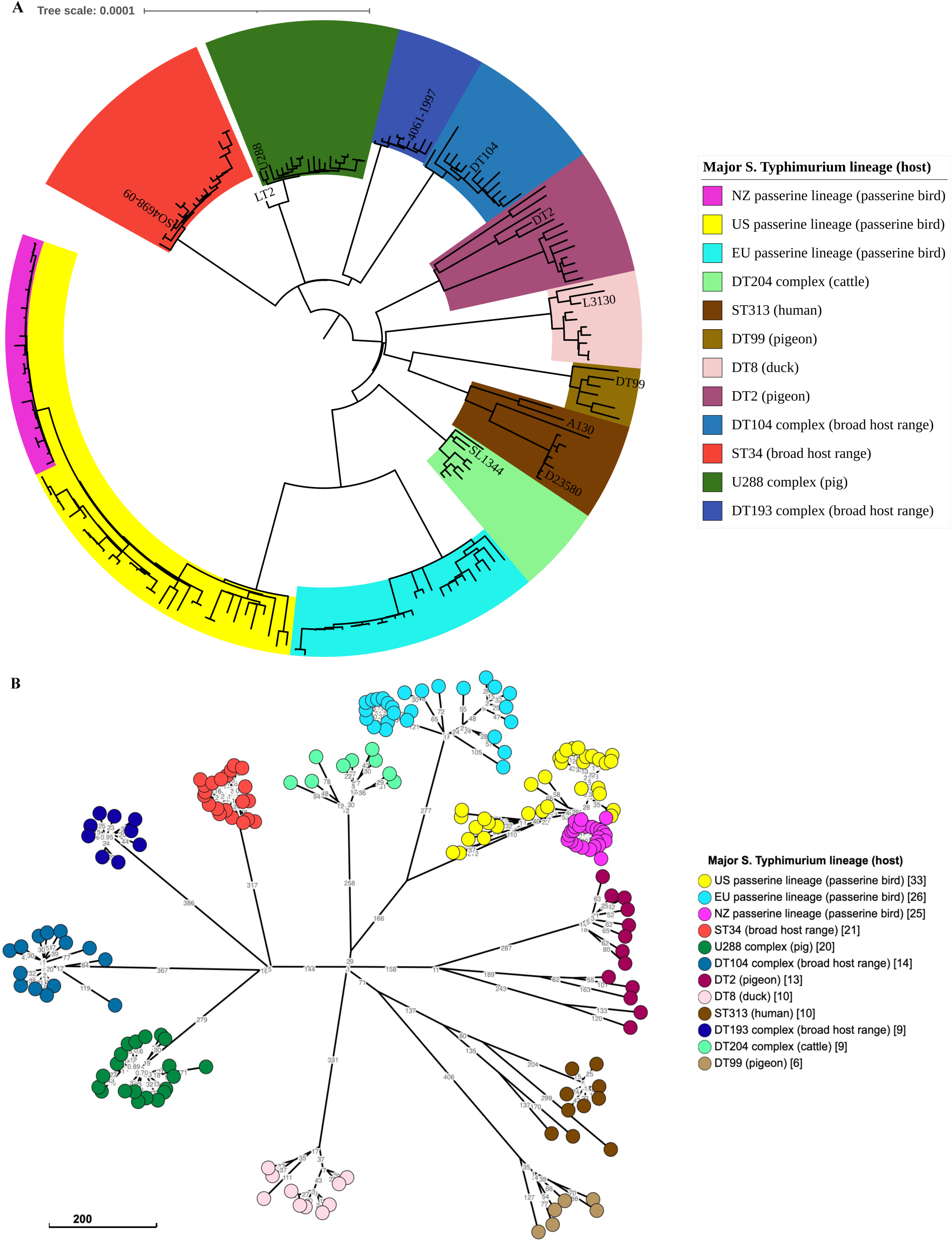
(**A**) Maximum-likelihood phylogenetic tree of the 196 *Salmonella enterica* serovar Typhimurium isolates from various hosts representing the genetic diversity within the serovar. The tree is created based on 10,065 core-genome single nucleotide polymorphisms (SNPs) with reference to *S*. Typhimurium LT2 and rooted at midpoint. Color ranges in the tree represent the major *S*. Typhimurium lineages identified in the literature and in this study. Labels at the tree tips represent the representative isolates from individual lineages. The legend field at the right of the tree represents the *S*. Typhimurium lineage (primary host). Broad host range in parentheses indicates that isolates from the corresponding lineage are commonly identified among humans, cattle, pigs, poultry, and other hosts or environmental niches. The specific host in parentheses indicates that isolates from the corresponding lineage are primarily from that specific host. (**B**) Neighbor joining tree of the 196 *S*. Typhimurium isolates from various hosts. The tree is created based on allelic differences in the 21,065 loci of the whole-genome multilocus sequence typing (wgMLST) *Salmonella* scheme with GrapeTree at EnteroBase. The major *S*. Typhimurium lineages are highlighted in colors in the tree. The legend field at the right of the tree represents the *S*. Typhimurium lineage (primary host) [number of isolates in the lineage]. The scale bar indicates 200 wgMLST alleles. Allele differences between isolates are indicated by numbers on the connecting lines.

### Genetic comparison of *S*. Typhimurium from different lineages

The average number of virulence genes, plasmid replicons, and AMR genes per isolate from a specific *S*. Typhimurium lineage is shown in Figure 4. Isolates from most of the individual *S*. Typhimurium lineages had an average number of 115–116 virulence genes. However, the average number of virulence genes per isolate from the EU, US passerine lineages and ST34 lineage was less than 110 (Figure 4). Similarly, isolates from these three lineages carried fewer plasmid replicons (average number <1) compared to isolates from other lineages (average number >1). In fact, we identified the absent virulence genes were mostly located on pSLT (*i.e*., *pefABCD, rck*, and *spvBCR*) (Table 3). Moreover, all of the isolates from the EU, NZ, US passerine lineages and DT99 lineage (host: pigeon) lacked identifiable AMR genes (average number = 1; the only AMR gene was *aac(6’)-Iaa*) (Figure 4). However, isolates from lineages with broad host range (average number >4), adapted to humans (ST313: average number ≈ 7), or associated with specific livestock (DT204 complex: average number ≈ 2; U288 complex: average number ≈ 8) had more AMR genes (Figure 4).

**Table 3.**
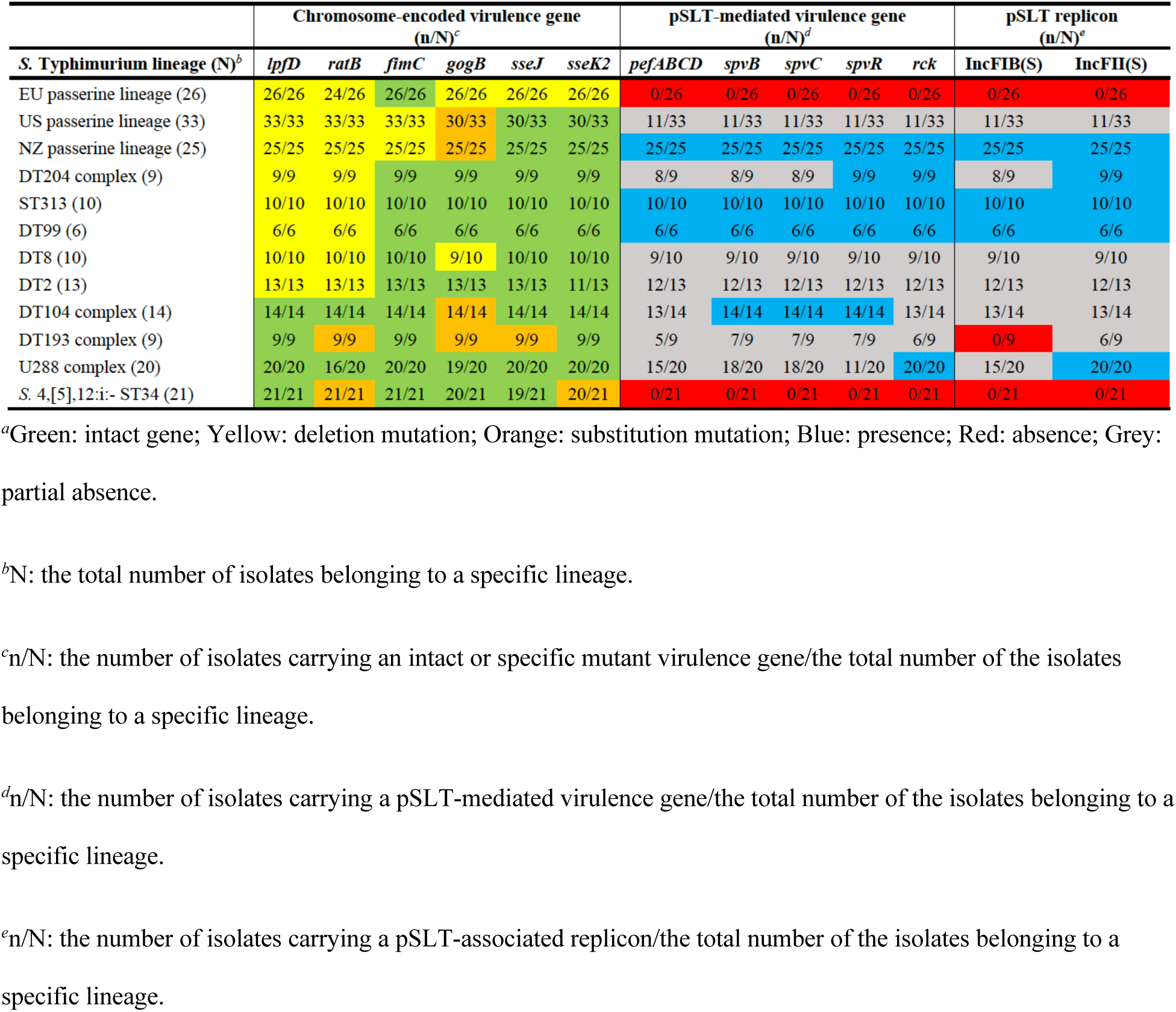
Differences in chromosome- and plasmid-mediated virulence genes between *Salmonella enterica* serovar Typhimurium isolates from diverse lineages^*a*^.

| *S. Typhimurium* lineage (N)<sup>b</sup> | Chromosome-encoded virulence gene<br>(n/N)<sup>c</sup> |  |  |  |  |  | pSLT-mediated virulence gene<br>(n/N)<sup>d</sup> |  |  |  |  | pSLT replicon<br>(n/N)<sup>e</sup> |  |
| --- | --- | --- | --- | --- | --- | --- | --- | --- | --- | --- | --- | --- | --- |
|  | *lpfD* | *ratB* | *fimC* | *gogB* | *sseJ* | *sseK2* | *pefABCD* | *spvB* | *spvC* | *spvR* | *rck* | IncFIB(S) | IncFII(S) |
| EU passerine lineage (26) | 26/26 | 24/26 | 26/26 | 26/26 | 26/26 | 26/26 | 0/26 | 0/26 | 0/26 | 0/26 | 0/26 | 0/26 | 0/26 |
| US passerine lineage (33) | 33/33 | 33/33 | 33/33 | 30/33 | 30/33 | 30/33 | 11/33 | 11/33 | 11/33 | 11/33 | 11/33 | 11/33 | 11/33 |
| NZ passerine lineage (25) | 25/25 | 25/25 | 25/25 | 25/25 | 25/25 | 25/25 | 25/25 | 25/25 | 25/25 | 25/25 | 25/25 | 25/25 | 25/25 |
| DT204 complex (9) | 9/9 | 9/9 | 9/9 | 9/9 | 9/9 | 9/9 | 8/9 | 8/9 | 8/9 | 9/9 | 9/9 | 8/9 | 9/9 |
| ST313 (10) | 10/10 | 10/10 | 10/10 | 10/10 | 10/10 | 10/10 | 10/10 | 10/10 | 10/10 | 10/10 | 10/10 | 10/10 | 10/10 |
| DT99 (6) | 6/6 | 6/6 | 6/6 | 6/6 | 6/6 | 6/6 | 6/6 | 6/6 | 6/6 | 6/6 | 6/6 | 6/6 | 6/6 |
| DT8 (10) | 10/10 | 10/10 | 10/10 | 9/10 | 10/10 | 10/10 | 9/10 | 9/10 | 9/10 | 9/10 | 9/10 | 9/10 | 9/10 |
| DT2 (13) | 13/13 | 13/13 | 13/13 | 13/13 | 13/13 | 11/13 | 12/13 | 12/13 | 12/13 | 12/13 | 12/13 | 12/13 | 12/13 |
| DT104 complex (14) | 14/14 | 14/14 | 14/14 | 14/14 | 14/14 | 14/14 | 13/14 | 14/14 | 14/14 | 14/14 | 13/14 | 13/14 | 13/14 |
| DT193 complex (9) | 9/9 | 9/9 | 9/9 | 9/9 | 9/9 | 9/9 | 5/9 | 7/9 | 7/9 | 7/9 | 6/9 | 0/9 | 6/9 |
| U288 complex (20) | 20/20 | 16/20 | 20/20 | 19/20 | 20/20 | 20/20 | 15/20 | 18/20 | 18/20 | 11/20 | 20/20 | 15/20 | 20/20 |
| *S. 4,[5],12:i:-* ST34 (21) | 21/21 | 21/21 | 21/21 | 20/21 | 19/21 | 20/21 | 0/21 | 0/21 | 0/21 | 0/21 | 0/21 | 0/21 | 0/21 |

<sup>a</sup>Green: intact gene; Yellow: deletion mutation; Orange: substitution mutation; Blue: presence; Red: absence; Grey: partial absence.
<sup>b</sup>N: the total number of isolates belonging to a specific lineage.
<sup>c</sup>n/N: the number of isolates carrying an intact or specific mutant virulence gene/the total number of the isolates belonging to a specific lineage.
<sup>d</sup>n/N: the number of isolates carrying a pSLT-mediated virulence gene/the total number of the isolates belonging to a specific lineage.
<sup>e</sup>n/N: the number of isolates carrying a pSLT-associated replicon/the total number of the isolates belonging to a specific lineage.

**Figure 4.**
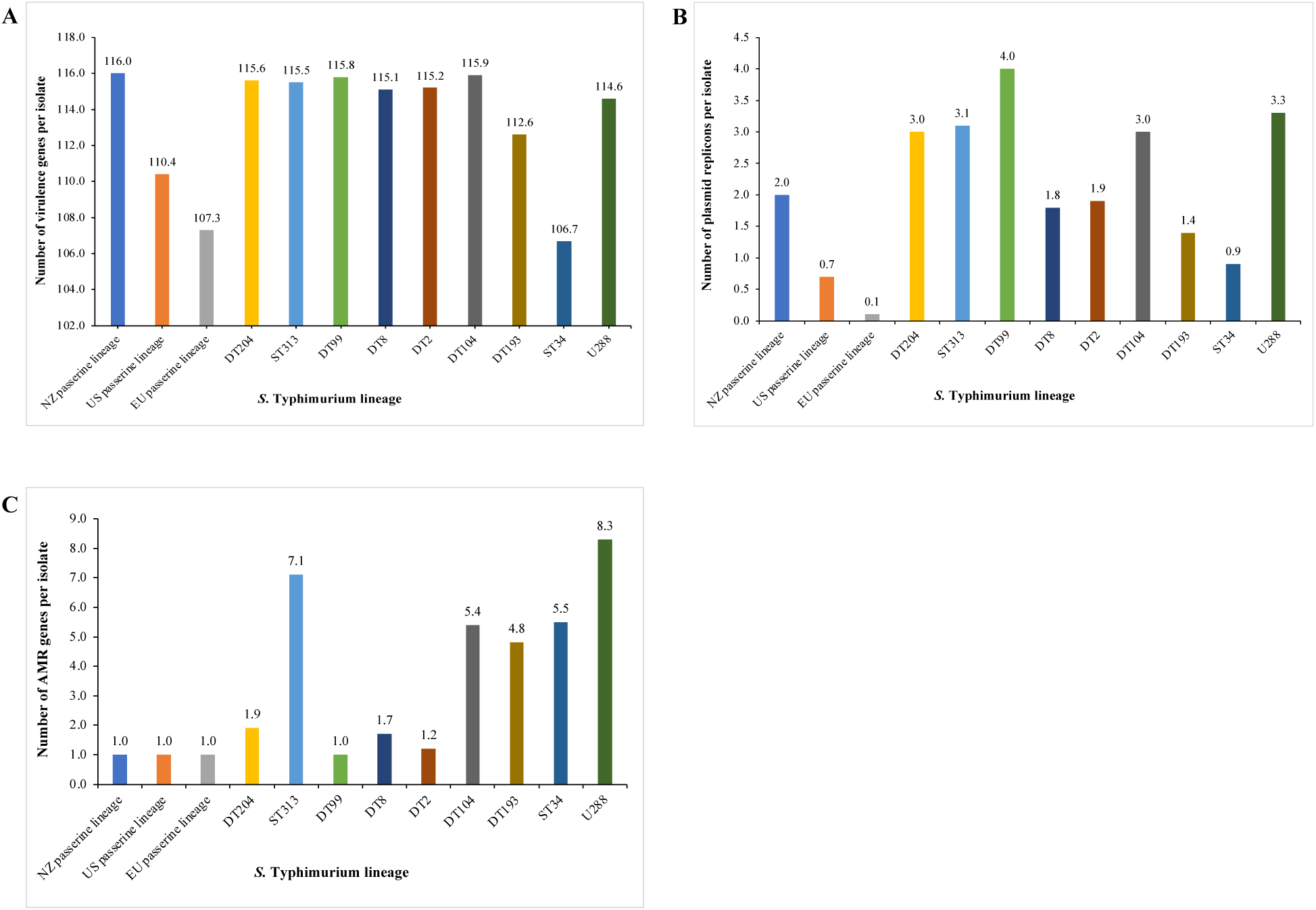
(**A**) Number of virulence genes per isolate detected in a specific *Salmonella enterica* serovar Typhimurium lineage. (**B**) Number of plasmid replicons per isolate detected in a specific *S*. Typhimurium lineage. (**C**) Number of antimicrobial resistance(AMR) genes per isolate detected in a specific *S*. Typhimurium lineage.

We also identified the virulence gene signatures that can discriminate passerine isolates from isolates of other hosts. Compared to isolates from other lineages, pseudogenization of *fimC* (full length: 708 bp; deletion of G at position 87) was unique to isolates from the US and NZ passerine lineages, while pseudogenization of *gogB* (full length: 1,498 bp; deletion of T at position 1,125), *sseJ* (full length: 1,234 bp; deletion of C at position 976), and *sseK2* (full length: 1,047 bp; deletion of A at position 522) only concurred in isolates from the EU passerine lineages (Table 3).

## DISCUSSION

Passerine-associated *S*. Typhimurium have a global distribution and are linked to human salmonellosis outbreaks in recent decades. In this study, we explored the emergence, genetic relationship, and evolution of passerine-associated *S*. Typhimurium from Europe, New Zealand, and the United States, and compared the passerine isolates with isolates from diverse hosts. Our study revealed that the EU and US passerine isolates formed two distinct lineages, while the NZ passerine isolates clustered as a sublineage of the US passerine lineage. Further, the emergence of the EU, NZ, and US passerine lineages were relatively recent events. Although the EU and US passerine lineages identified in this study were distinct from each other with different virulence genetic signatures, they clustered in a lineage that was distantly related to the major *S*. Typhimurium lineages formed by isolates from multiple hosts (*i.e*., humans, cattle, pigs, poultry, pigeons, ducks). One of the caveats of the study is a general paucity of whole-genome sequences of passerine isolates from Asia, Africa, South America in public database (*e.g*., NCBI). Integration of passerine isolates from these locations will improve the robustness of the genomic analysis.

Previous epidemiologic survey indicated that passerine salmonellosis was caused by specific *S*. Typhimurium variants. For examples, *S*. Typhimurium DT40 and DT56(v) were the dominant *S*. Typhimurium isolated from passerines suffering from salmonellosis in European countries such as the United Kingdom and Sweden (5, 9, 10). Additionally, *S*. Typhimurium DT160 were demonstrated to be responsible for the human and passerine salmonellosis outbreaks in New Zealand and Australia (7, 8). In the United States, *S*. Typhimurium PFGE (pulsed-field gel electrophoresis) type A3 from pine siskins was identified as the cause of salmonellosis outbreaks both in humans and passerines (11). Although these epidemiologic studies were able to link specific passerine-associated *S*. Typhimurium variants to human and passerine salmonellosis outbreaks through phage typing or PFGE typing, the genetic relationship of these pathovariants from different locations remains unknown. The availability of whole-genome sequences through public database facilitates a high-resolution investigation on the genetic relatedness of globally sourced passerine isolates.

The 84 passerine isolates from the United Kingdom, Sweden, Germany, New Zealand, and the United States clustered in a lineage distinct from the *S*. Typhimurium lineages that have broad host range (*e.g*., humans, livestock, poultry), such as DT104 complex (1), monophasic *S*. Typhimurium ST34 (2), and DT193 complex (14). The passerine lineage also differed from those narrow-host-range lineages such as U288 complex lineage primarily circulating in pigs (15), DT204 complex lineage majorly infecting cattle (16), DT2 (17) and DT99 (3) lineages adapted to pigeons, DT8 lineage adapted to ducks (4), and ST313 lineage (18, 19) causing invasive human salmonellosis in sub-Saharan Africa. The distinct phylogenetic lineage formed by geographically dispersed passerine isolates agrees with a previous study, in which Mather et al. (2016) demonstrated that the UK passerine isolates formed a lineage distinct from representative isolates of diverse hosts (*e.g*., humans, cattle, horses, chicken, pigeons) and geographical regions (20). The phylogenetic signature presented by the passerine isolates from different countries supported the hypothesis that certain *S*. Typhimurium variants have undergone evolution towards a passerine-adapted lifestyle.

Host adaptation is often accompanied by loss-of-function mutation or genome degradation (18, 21–23). For example, loss of virulence for secondary hosts have been observed in *S*. Typhimurium adapted to pigeons (17). In this study, pseudogenization of *lpfD* and *ratB* due to deletion mutation was found in host-adapted *S*. Typhimurium isolates from the DT2, DT8, DT99, DT204 complex, ST313, and the passerine lineages, except isolates from U288 complex lineage possibly adapted to pigs (Table 3). Therefore, the two virulence genes may segregate host-adapted *S*. Typhimurium lineages from lineages with broad host range (*i.e*., ST34, DT104 complex, DT193 complex). In addition, the passerine isolates accumulated virulence pseudogenes other than *lpfD* and *ratB*. The US and NZ passerine isolates had a single-base deletion in type 1 fimbrial gene *fimC*, while the EU passerine isolates had a concurrent single-base deletion in T3SS effector genes *gogB, sseJ*, and *sseK2*. It should be noted these virulence genes were intact in almost all of the isolates from other lineages (Table 3). Therefore, they may be genetic signatures that can discriminate passerine isolates from other-sourced isolates. Further, the *fimC* gene is required for the biosynthesis of type 1 fimbriae, which is involved in adhesion to host cells (24). It has been reported that allelic variation in type 1 fimbriae affected *Salmonella* host specificity (22). In addition, a recent study reported that loss-of-function mutations in T3SS effector genes attenuated pathogenicity of *S*. Typhimurium to humans or mammals but maintained virulence to avian hosts (25). Taken together, pseudogenization of fimbrial or T3SS effector genes due to frameshift mutation may lead to loss of virulence and contribute to host adaptation of *S*. Typhimurium to the passerine hosts.

The plasmid pSLT is an *S*. Typhimurium-specific virulence plasmid that harbors virulence genes such as *pefABCD, rck*, and *spvBCR* (26). These virulence genes are important for *S*. Typhimurium survival and replication in human and mouse macrophages (25, 27, 28). All of the EU passerine isolates (26/26) and two thirds (22/33) of the US passerine isolates lacked pSLT, while most of the isolates from other lineages except monophasic *S*. Typhimurium ST34 harbored this virulence plasmid (Table 3), indicating that the pSLT-mediated virulence was dispensable for *S*. Typhimurium pathogenesis in passerine hosts. The loss of pSLT in passerine isolates is likely an ongoing process as pSLT is only absent in a partial number of the US and NZ passerine isolates. In addition to lack of pSLT, the EU, US, and NZ passerine isolates also lacked AMR genes with the exception of *aac(6’)-Iaa*. The absence of AMR genes occurred in DT99 isolates from feral pigeons as well. In contrast, AMR genes were identified in isolates from other diverse hosts, especially those with broad host range (Figure 4). The lack of AMR genes in globally distributed passerine isolates is consistent with our previous study, which revealed the low occurrence of AMR in *S*. Typhimurium from wild birds in the United States (29). A plausible explanation for the observation is that environments utilized by passerine birds are less exposed to antibiotics compared to those utilized by domestic animals and humans. Therefore, isolates from passerines are rarely subjected to antibiotic selection pressure and thus less likely to develop AMR.

The passerine isolates from European countries and the United States formed two lineages closely related to each other (Figure 3; average SNP distance in the core genome ≈ 265), suggesting common ancestry for the two lineages (30). Further, Bayesian inference suggested that the MRCA of the two lineages originated from *ca*. 1840 (Figure 2). As the EU and US passerine lineages were more closely related (average SNP distance in the core genome ≈ 308) to DT204 complex lineage (primary host: cattle) compared to other lineages in the tree (Figure 3), it is possible that passerine birds acquired the MRCA from domestic animals, potentially cattle. However, more evidence is required to determine the original host of the MRCA and the directionality of transmission. In addition, the emergence of the US, EU, and NZ passerine lineages were estimated after 1950 over short timescales, indicating that host adaptation of *S*. Typhimurium to passerines may be a relatively recent ongoing process driven by anthropogenic influences. The passerine isolates (*i.e*., *S*. Typhimurium DT160) from New Zealand clustered as a sublineage of the US passerine lineage.

Further, the NZ passerine lineage was estimated to emerge in *ca*. 1995–1997 in this study. In a previous study, Bloomfield et al. (2017) reported that the salmonellosis outbreak caused by *S*. Typhimurium DT160 resulted from a single introduction into New Zealand between 1996 and 1998 (7). However, the origin and source of the outbreak have not been identified. Our study reveals the closely genetic relatedness (average SNP distance in the core genome ≈ 81) between isolates from the NZ and US passerine lineages, suggesting that DT160 isolates in New Zealand may have originated from passerines outside Oceania. However, whole-genome sequences of passerine isolates from other locations (*i.e*., Asia, South America, and Africa) are necessary to conduct further genomic analysis to test this hypothesis.

In conclusion, our study demonstrates the importance of whole-genome sequencing and genomic analysis of historical microbial collections. The findings provide insights into host adaptation of *S*. Typhimurium in passerines and are helpful for modern day epidemiologic surveillance. Host-specific genetic signatures identified in this study can aid source attribution of *S*. Typhimurium to avian hosts in outbreak investigation. As passerines are highly mobile and can spread zoonotic pathogens over a large spatial scale, it is important to raise our awareness of passerines as reservoirs of specific *S*. Typhimurium variants. Although these pathovariants only account for a small number of human salmonellosis cases worldwide, control strategies, for example washing hands after contact with wild birds, would be taken to reduce potential transmission between passerines and humans.

## MATERIALS AND METHODS

### Dataset selection and quality assessment for raw reads

Passerine-associated *S*. Typhimurium isolates (Table 1; New Zealand: *n* = 25, isolated year: 2000–2009; United States: *n* = 33, isolated year: 1978–2019; European countries: *n* = 26, isolated year: 2001–2016) were derived from wild birds with confirmed salmonellosis over broad temporal and spatial scales, and some of these isolates also had closely genetic relatedness with human clinical isolates (7, 10, 20, 31). Therefore, the isolates were chosen to represent *S*. Typhimurium pathovariants emerging worldwide that caused salmonellosis both in humans and passerines. Context *S*. Typhimurium isolates (Dataset S2; *n* = 112) were selected to represent the phylogenetic diversity of this serovar across different hosts and geographic locations, and to compare the genomic differences between isolates from passerines and other multiple hosts. The Illumina paired-end reads of the selected isolates were available at the NCBI database (accession number provided in Table 1 and Dataset S2). The quality of the sequence data was assessed using the MicroRunQC workflow in GalaxyTrakr v2 (32). Raw reads meeting the quality control requirements (*i.e*., average coverage >30, average quality score >30, number of contigs <400, total assembly length between 4.4–5.l Mb) were used for genomic analysis in this study.

### Phylogenetic analysis

The phylogenetic relationship of the 84 passerine isolates was inferred from their core genomes. Snippy (Galaxy v4.5.0) (https://github.com/tseemann/snippy) was used to generate a whole-genome alignment and find SNPs between the reference genome LT2 (RefSeq NC_003197.1) and the genomes of passerine isolates. Snippy-core (Galaxy v4.5.0) (https://github.com/tseemann/snippy) was used to convert the Snippy outputs (*i.e*., whole-genome alignment) into a core-genome alignment. The resultant core-genome alignment (2,253 SNPs in the core genomic regions) was used to construct a maximum-likelihood phylogenetic tree by MEGA X (v10.1.8) (33) using the Tamura-Nei model and 500 bootstrap replicates. The SNP phylogenetic tree was visualized and annotated using the Interactive Tree of Life (iTOL v6; https://itol.embl.de). SNP distance between sequences was calculated using snp-dists (Galaxy v0.6.3) (https://github.com/tseemann/snp-dists). Sequence type (ST) of the *S*. Typhimurium isolates was identified using 7-gene (*aroC, dnaN, hemD, hisD, purE, sucA*, and *thrA*) MLST at EnteroBase (34). STs were then annotated in the SNP phylogenetic tree. We also generated a maximum-likelihood phylogenetic tree of the 84 passerine isolates (Table 1) and 112 context isolates (Dataset S2) from diverse hosts to represent the genetic diversity within serovar Typhimurium. The tree was created based on 10,065 SNPs in the core genomic regions of the 196 passerine and context isolates with reference to *S*. Typhimurium LT2 using the EnteroBase SNP project (34). In addition, a NJ tree of the 196 passerine and context isolates based on the *Salmonella* whole-genome MLST (21,065 loci) scheme at EnteroBase (34) was built to complement the core-genome SNP-based phylogenetic analysis.

### Bayesian inference

A time-scaled Bayesian phylogenetic tree was constructed to determine the divergence times of the *S*. Typhimurium lineages from passerines. The temporal signal of the sequence data was examined using TempEst (35) before phylogenetic molecular clock analysis (Figure S1). The core-genome alignment (2,253 SNPs in the core genomic regions) of passerine isolates (*n* = 84; Table 1) generated previously was used as the input for the time-scaled tree construction. The parameters for constructing the Bayesian phylogenetic tree were set in BEAUti (v2.6.5) (36) as follows: Prior assumption-coalescent Bayesian skyline; Clock model-relaxed clock log normal with the default clock rate value of 1.0; and Markov chain Monte Carlo (MCMC) at chain length-100 million, storing every 1,000 generations. Two independent runs with the same parameters were performed in BEAST2 (v2.6.5) (36) to ensure convergence. The resultant log files were viewed in Tracer (v1.7.2) to check if the effective sample sizes of all parameters were more than 200 and the MCMC chains were converged. A maximum clade credibility tree was created using TreeAnnotator (v2.6.4) (36) with a burn-in percentage of 10% and node option of median height. Finally, the tree was visualized using FigTree v1.4.4 (https://github.com/rambaut/figtree/releases). To determine the substitution rate for the genome of passerine isolates, we multiplied the substitution rate estimated by BEAST2 (v2.6.5) by the number of analyzed core-genome SNPs (2,253 bp), and then divided the product by the average genome size of the analyzed passerine isolates (4,951,383 bp).

### Antimicrobial resistance, virulence, and plasmid profiling

Raw reads of each isolate were *de novo* assembled using Shovill (Galaxy v1.0.4) (37). ABRicate (Galaxy v1.0.1) (38) was used to identify the AMR genes, virulence factors, and plasmid replicons by aligning each draft genome assembly against the ResFinder database (39), VFDB (40), and PlasmidFinder database (41), respectively. For all searches using ABRicate, minimum nucleotide identity and coverage thresholds of 80% and 80% were used, respectively. Virulence genes that were not 100% identical or covered with the reference virulence gene from VFDB may have deletions, insertions, or substitutions of interest. We then manually checked the mutation type by aligning the virulence gene of interest against the reference virulence gene from VFDB using BLAST (https://blast.ncbi.nlm.nih.gov/Blast.cgi).

### Data availability

Sequence data of the *S*. Typhimurium strains are publicly available in the NCBI Sequence Read Archive (https://www.ncbi.nlm.nih.gov/sra). Accession numbers are available in Table 1 and Dataset S2.

## ACKNOWLEDGMENTS

This work is supported by the Food and Drug Administration (US Department of Health and Human Services) (Grant No. 1U19FD007114-01), US Department of Agriculture (Grant No. PEN4522), and Penn State College of Agricultural Sciences. It is also supported in part by the Food and Drug Administration (Grant No. 1U01FD006253-01 to Pennsylvania Department of Health for collaboration in the National Antimicrobial Resistance Monitoring System).

## AUTHOR CONTRIBUTION

Y.F. designed the study, sequenced the US passerine isolates, collected the globally sourced data from EnteroBase and NCBI, performed the comparative genomic analysis of the data, interpreted the data, and wrote the draft manuscript; N.M.M. and E.G.D. contributed to interpretation of the data and manuscript revision.

## CONFLICTS OF INTEREST

The authors declare no competing interests.

---

**Figure S1.**
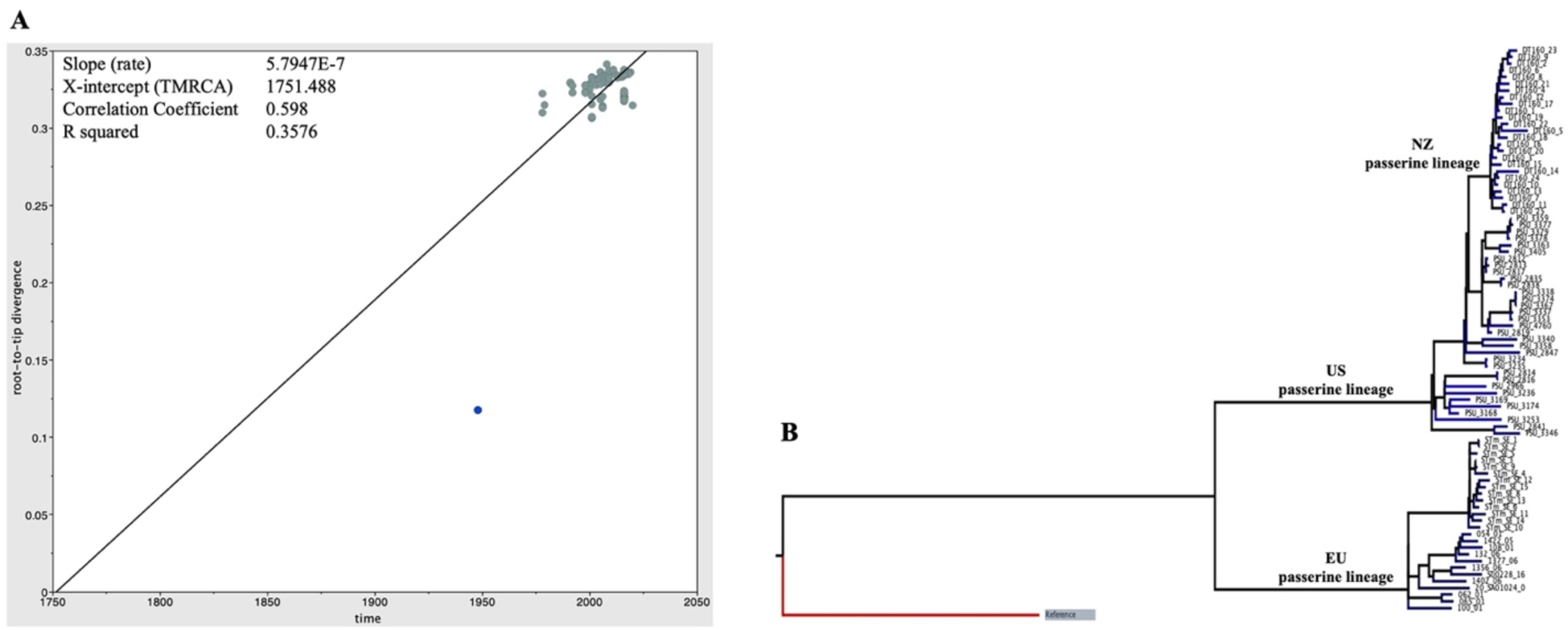
Temporal signal of the 84 *S*. Typhimurium genome sequences from passerine birds used for Bayesian inference. (**A**) Root-to-tip regression plot showing regression of genetic distance against sampling time. (**B**) Phylogeny of 84 *S*. Typhimurium genome sequences from passerine birds. The EU, US, and NZ passerine lineages are indicated on the phylogenetic tree branches. Reference genome from *S*. Typhimurium LT2 is highlighted in blue in (**A**) and shaded in grey in (**B**).

## Supplementary Dataset Legend

**Dataset S1**. *In silico* virulence, antimicrobial resistance (AMR), and plasmid profiles of the public available *Salmonella enterica* serovar Typhimurium isolates from passerine birds (*n* = 84) and other diverse hosts (*n* = 112).

**Dataset S2**. Metadata information of the 112 context *Salmonella enterica* serovar Typhimurium isolates from diverse hosts.

## Notes

### Competing Interest Statement

The authors have declared no competing interest.

